# Theory of active chromatin remodeling

**DOI:** 10.1101/687145

**Authors:** Zhongling Jiang, Bin Zhang

## Abstract

Nucleosome positioning controls the accessible regions of chromatin and plays essential roles in DNA-templated processes. ATP driven remodeling enzymes are known to be crucial for its establishment *in vivo*, but their non-equilibrium nature has hindered the development of a unified theoretical framework for nucleosome positioning. Using a perturbation theory, we show that the effect of these enzymes can be well approximated by effective equilibrium models with rescaled temperatures and interactions. Numerical simulations support the accuracy of the theory in predicting both kinetic and steady-state quantities, including the effective temperature and the radial distribution function, in biologically relevant regimes. The energy landscape view emerging from our study provides an intuitive understanding for the impact of remodeling enzymes in either reinforcing or overwriting intrinsic signals for nucleosome positioning, and may help improve the accuracy of computational models for its prediction in silico.

---

The eukaryotic genome is packaged into nucleosomes that wrap approximately 147-base-pair (bp) stretch of DNA around histone proteins [1]. Formation of nucleosomes helps to fit the genome inside the nucleus, but also occludes the binding of protein molecules to the DNA [2, 3]. The precise position of nucleosomes along the chromatin, therefore, is of critical importance for gene regulation as it determines which part of the genome is accessible for binding by regulatory factors and transcriptional machinery [4, 5]. Remarkably, nucleosomes do not uniformly cover the whole genome but favor specific positions, and are depleted at many promoter and enhancer regions to accommodate transcription. Underpinning the molecular determinants of nucleosome positioning is of fundamental interest and can provide insight into the regulation of gene expressions [6–8].

Nucleosome positioning *in vivo* is intimately related to the underlying DNA sequence that dictates the bendability of the DNA molecule and the stability of formed nucleosomes [9]. Genome-wide studies have found a ~ 10-bp periodicity of bendable dinucleotides (AT and TA) along the nucleosome length [6, 10, 11], and that many nucleosome-depleted regions are enriched with intrinsically stiff poly(dA:dT) tracts [12]. Computational models based on such sequence features have achieved a varied degree of success in predicting nucleosome positioning [13]. These models are often of thermodynamic nature and, therefore, inherently limited due to their neglect of chromatin remodeling enzymes that drive the system out of equilibrium [14, 15]. Though the impact of chromatin remodelers has been demonstrated [16, 17], conflicting interpretations have been reported for them to either reinforce or overwrite positioning signals from the DNA sequence. Theoretical studies on remodeling enzymes are challenging due to the non-equilibrium nature and have been mostly limited to kinetic simulations [18–20]. Here, we develop an analytical theory to provide a unified framework for studying the effect of remodeling enzymes, and to reconcile their combined role with intrinsic thermodynamic signals in positioning nucleosomes. Our study also provides a theoretical foundation for using effective equilibrium models to analyze experimental data [21, 22], and may help to develop quantitative models that incorporate remodeling effects to predict nucleosome positioning. In the following, we first introduce a simplified kinetic model for nucleosome positioning that explicitly considers the effect of two different remodeling enzymes. We then show that kinetic and steady state properties of this non-equilibrium system can be rigorously mapped onto an effective equilibrium model. Finally, results from numerical simulations are provided to validate the accuracy of the theory.

We used a one-dimensional lattice model to study nucleosome positioning, with each discrete site corresponding to one base pair of DNA (see Figure 1). The position of a nucleosome *i* is noted with *x*_*i*_, which corresponds to the location of the nucleosome center. A soft-core potential *v*(Δ*x*) was applied between neighboring nucleosomes separated by Δ*x* base pairs to account for the excluded volume effect, as well as transient DNA unwrapping [22].

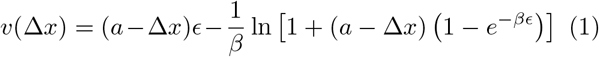

if Δ*x* < *a* and 0 otherwise. *ϵ* = 0.1525 *k*_*B*_*T* measures the energetic cost of unwinding one bp and *β* = 1/*k*_*B*_*T*. In contrast to a hard-core potential that would strictly enforce a minimum separation, *v*(Δ*x*) allows for significant overlap between neighboring nucleosomes at a finite energetic cost. To focus on the role of remodeling enzymes, we did not explicitly model the DNA sequence effect, and assume that all lattice sites share the same binding affinity. The theory presented below, however, can be easily generalized to include DNA specificity.

For simplicity, we considered systems with fixed nucleosome density and did not include nucleosome formation and disassembly processes. The kinetics of our model consists of only diffusion and active remodeling (see Figure 1). The diffusion rate *d*_*mn*_ for a nucleosome to move from site *m* to *n* satisfies detailed balance and 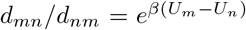, where *U*_*m*/*n*_ = Σ_*i*_*v*(Δ*x*_*i*_ = *x*_*i*+1_ − *x*_*i*_) is the total energy of the system before and after movement, respectively. Following Padinhateeri and Marko [18], we define 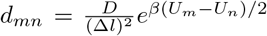, with *D* being the diffusion coefficient, and Δ*l* = |*m* − *n*|. Numerous enzymes have been discovered to organize nucleosomes along the DNA by consuming ATP, and we focus on two representative ones with distinct kinetics. Type one enzymes interact with a single nucleosome, and can move it to left or right by *l* base pairs with a rate of *k*_1_. These enzymes resemble the function of SWI/SNF [23] and can significantly fluidize nucleosomes to achieve high packing density [18]. Type two enzymes, on the other hand, interact with a neighboring pair of nucleosomes, and can bring them closer to each other by *l* base pairs at a rate of *k*_2_ if the two are within Δ*x*_max_ = 332 bp [22]. A typical example for these enzymes is the ATP-utilizing chromatin assembly and remodeling factor (ACF) from the ISWI family, which is crucial for the regular spacing of nucleosomes near transcription start sites [24]. Since enzyme rates *k*_1/2_ are independent of the system’s energy, detailed balance is violated.

**FIG. 1.**
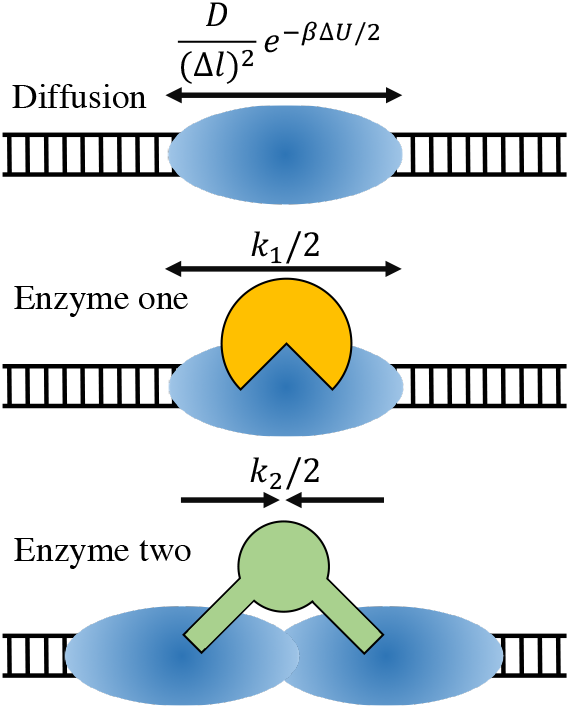
Illustration of the kinetic model for nucleosome positioning that includes diffusion (top), ATP-driven single-nucleosome remodeling (middle), and ATP-driven remodeling for a pair of nucleosomes (bottom). The DNA is drawn as a black ladder, with nucleosomes shown in blue oval and the two enzymes drawn in yellow and green respectively. The rates for an elementary step of different dynamics are shown above the arrows.

Our goal is to approximate both kinetic and steady-state properties of the above non-equilibrium system with effective equilibrium models. To make progress, we first describe the dynamical evolution of nucleosome positions **x** = {*x*_1_,*x*_2_,…,*x*_*n*_} with the following master equation

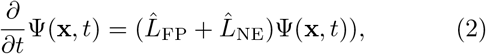

where Ψ(**x**,*t*) is the time dependent probability distribution function and 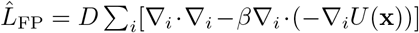 is the many-body Fokker-Planck operator describing the diffusive dynamics of nucleosomes. 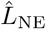 is the non-equilibrium operator for remodeling enzymes. For type one enzymes, it adopts the following expression

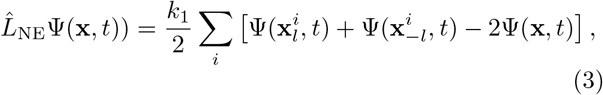

where 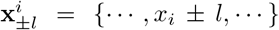. Similarly, the non-equilibrium operator for type two enzymes can be written as

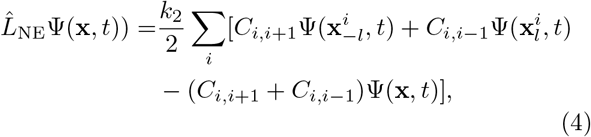

where *C*_*i,i*±1_ = 1 for 0 < |*x*_*i*_ − *x*_*i*±1_| ≤ Δ*x*_max_ and 0 otherwise. We note that similar equations have been used by Wolynes and coworkers to study the effect of molecular motors on the actin network [25].

Next, following Wang and Wolynes [26, 27], we expand the probability distribution function Ψ(**x***, t*)) in the powers of *l* up to the quadratic order. This leads to the following effective Fokker-Planck equation

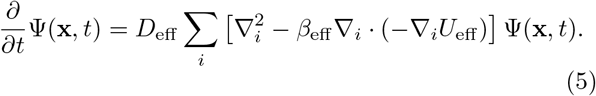

For Type one enzymes, we have

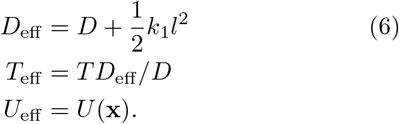

For Type two enzymes, we have

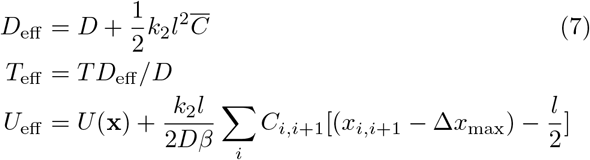

where 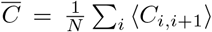, and 〈⋅〉 represents ensemble averaging. Eqs. 6 and 7 are the main results of this manuscript, and they provide intuitive interpretations for the role of remodeling enzymes in nucleosome positioning. Type one enzymes elevate the system’s temperature without perturbing the underlying energy landscape, while type two enzymes give rise to an attractive potential between neighboring nucleosomes in addition to a rescaled temperature.

Eq. 5 suggests that non-equilibrium systems can be rigorously mapped onto effective equilibrium models in the small *l* limit. To validate the accuracy of the derived effective models, we carried out stochastic simulations of the full kinetic model using the Gillespie algorithm [28] to obtain “exact” non-equilibrium results. A 58800 bp long DNA with a total of 400 binding sites was used together with the periodic boundary condition. Only one type of enzymes is included in any given simulation in order to study their effects separately. We used a nucleosome density of 0.8 for simulations with type one enzymes and 0.5 for type two enzymes. These densities were chosen to highlight the effect of the enzymes and do not affect our conclusions. More details of these simulations are provided in the Supporting Information (SI) [29].

We first determined the effective diffusion constants for type one enzymes from linear fitting of the mean-squared displacement 〈(*x*(*t*) − *x*(0))^2^〉 = 2*D*_eff_*t*. As shown in Figure 2(a), at a fixed rate *k*_1_ = 1 s^−1^, simulated values for *D*_eff_ (red dots) indeed closely matches the quadratic relationship with *l* as predicted in Eq. 6 (blue line). At large *l* values (*l* > 10 bp), the theory begins to deviate from numerical simulations. Upon close inspection of simulated configurations, we found significant overlap between neighboring nucleosomes at large *l* (see Figure S1). As chromatin remodeling is ignorant of the underlying energy landscape, performing enzymatic reactions on these strongly overlapping configurations can lead to nucleosomes passing through each other, which is prohibited in our simulation. As a result, many remodeling steps will be rejected, effectively causing a slow down of the diffusion. To support this argument, we performed additional short timescale simulations initialized with well separated nucleosomes. *D*_eff_ determined from these simulations are indeed in a good agreement with theoretical values over a much wider range (Figure S2). It’s worth noting that the biological value for *l* has been estimated to be 1 bp [30], well within the regime where the theory works. We further determined *D*_eff_ for various enzyme rates *k*_1_ at *l* = 1 bp, and again found excellent agreement between theory (purple line) and numerical simulations (green dots).

**FIG. 2.**
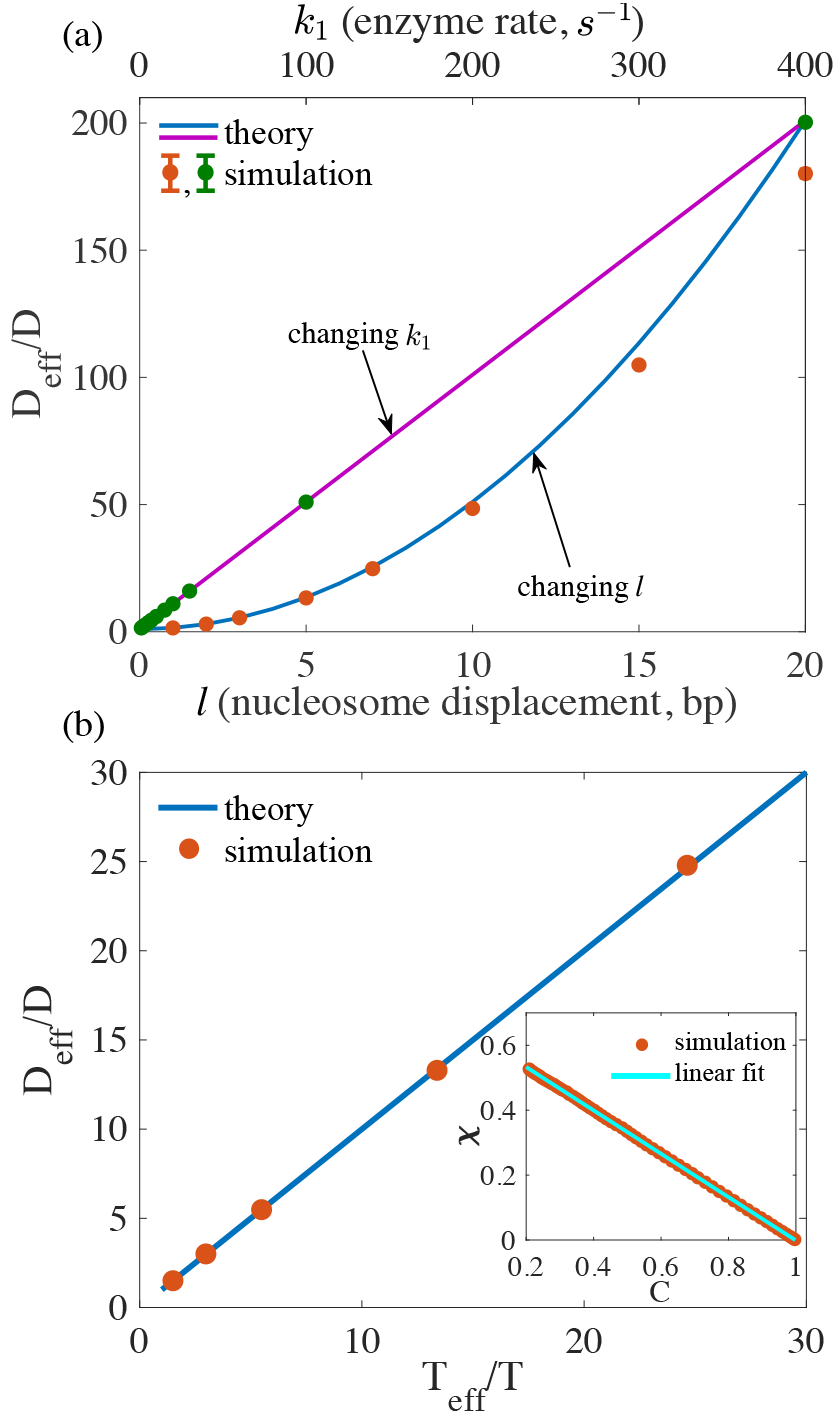
Comparison between simulated and theoretical values of *D*_eff_ and *T*_eff_ for type one enzymes. (a) Dependence of *D*_eff_ on the enzyme rate *k*_1_ and the nucleosome displacement *l* per enzymatic step. (b) *T*_eff_ determined as the fluctuation-dissipation ratio at *l* = 1, 2, 3, 5, 7 bp and *k*_1_ = 1 s^−1^. Error bars measured as standard deviation of the mean are comparable to the size of the symbols.

A notable result of the theory is that the fluctuation-dissipation relationship is satisfied with effective parameters. As a test, we determined the effective temperature using the fluctuation-dissipation ratio 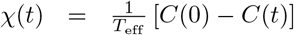 [31, 32]. We define *χ*(*t*) = 〈(*O*(*t*) − *O*(0)〉/*h*, *C*(*t*) = 〈(*O*(*t* + *t*_*o*_)*O′*(*t*_*o*_)〉 − 〈*O*(*t*_*o*_)〉 〈(*O′*(*t*_*o*_)〉, 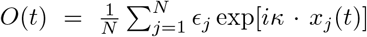 and 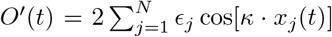. *ϵ*_*j*_ are random numbers with values of ±1 and the wave vector *κ* is chosen as the value for the first peak of the structure factor [29]. As shown in the inset of Figure 2(b), a linear relationship is found between *χ*(*t*) and *C*(*t*). In the main frame, we plotted *T*_eff_ for various values of *l*. The simulation again agrees well with theoretical prediction and *T*_eff_ and *D*_eff_ indeed follow a linear relationship.

To validate the accuracy of the effective equilibrium model in predicting steady state quantities, we computed the *g*(*r*) using nucleosome configurations obtained from kinetic simulations for type one enzymes with different step sizes. As shown in Figure 3 as empty circles, consistent with high effective temperatures, the systems with large step sizes gradually lose the structural ordering, and the peaks in *g*(*r*) disappear. The high probability at *r* close to 0 for *l* = 5 bp again suggests the presence of overlap between nucleosomes. We further calculated *g*(*r*) for effective equilibrium models (cyan lines) using a matrix treatment of the grand partition function proposed by Poland [33], and found that they are in excellent agreement with the ones calculated from kinetic simulations.

**FIG. 3.**
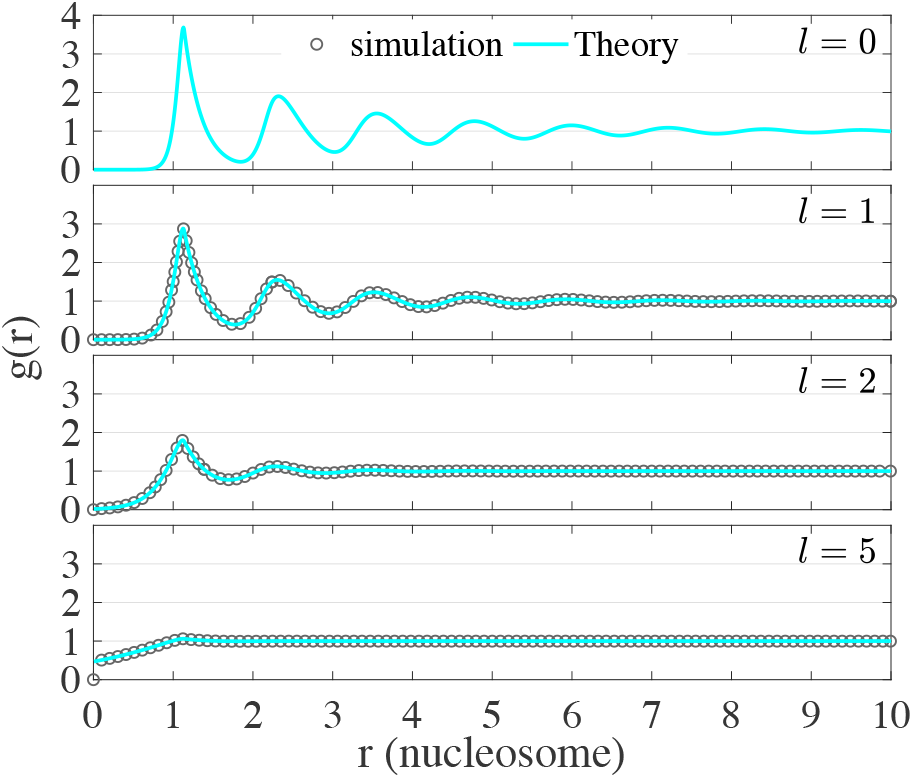
Comparison between radial distribution functions obtained from non-equilibrium simulations (empty circles) and from theoretical predictions of the effective equilibrium model (cyan lines) for type one enzymes with various step sizes.

The results for type one enzymes are quite encouraging and support the accuracy of the perturbation theory. We next study type two enzymes that have been shown to introduce structural ordering to the system, and produce chromatin segments with regular nucleosome spacing [22]. Our analytical expressions suggest that the ordering arises from an effective attraction between neighboring nucleosomes.

To validate the accuracy of the effective models, we again carried out non-equilibrium simulations with explicit enzymes using different step sizes *l*. As shown in Figure 4(a), we observed an increase of the diffusion coefficient at larger *l* due to contributions from ATP driven nucleosome sliding (red dots). The theoretical results (blue dots) were calculated using Eq. 7 with steady state values for 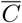 determined from non-equilibrium simulations. Though they agree well at small step sizes, theoretical results begin to deviate significantly from simulation values for *l >* 5 bp. Similar behaviors are seen for *D*_eff_ calculated at different enzyme remodeling rates *k*_2_ (purple and green dots). The difference between theory and simulation can again be attributed to the emergence of overlapping nucleosome configurations and rejection of remodeling steps that result in nucleosomes crossing each other (Figures S3 and S4). We further followed the same approach as type one enzymes to calculate the effective temperature from the fluctuation dissipation ratio. As shown in Fig. 4(b), *T*_eff_ again satisfies a linear relationship with *D*_eff_ as predicted by Eq. 7.

**FIG. 4.**
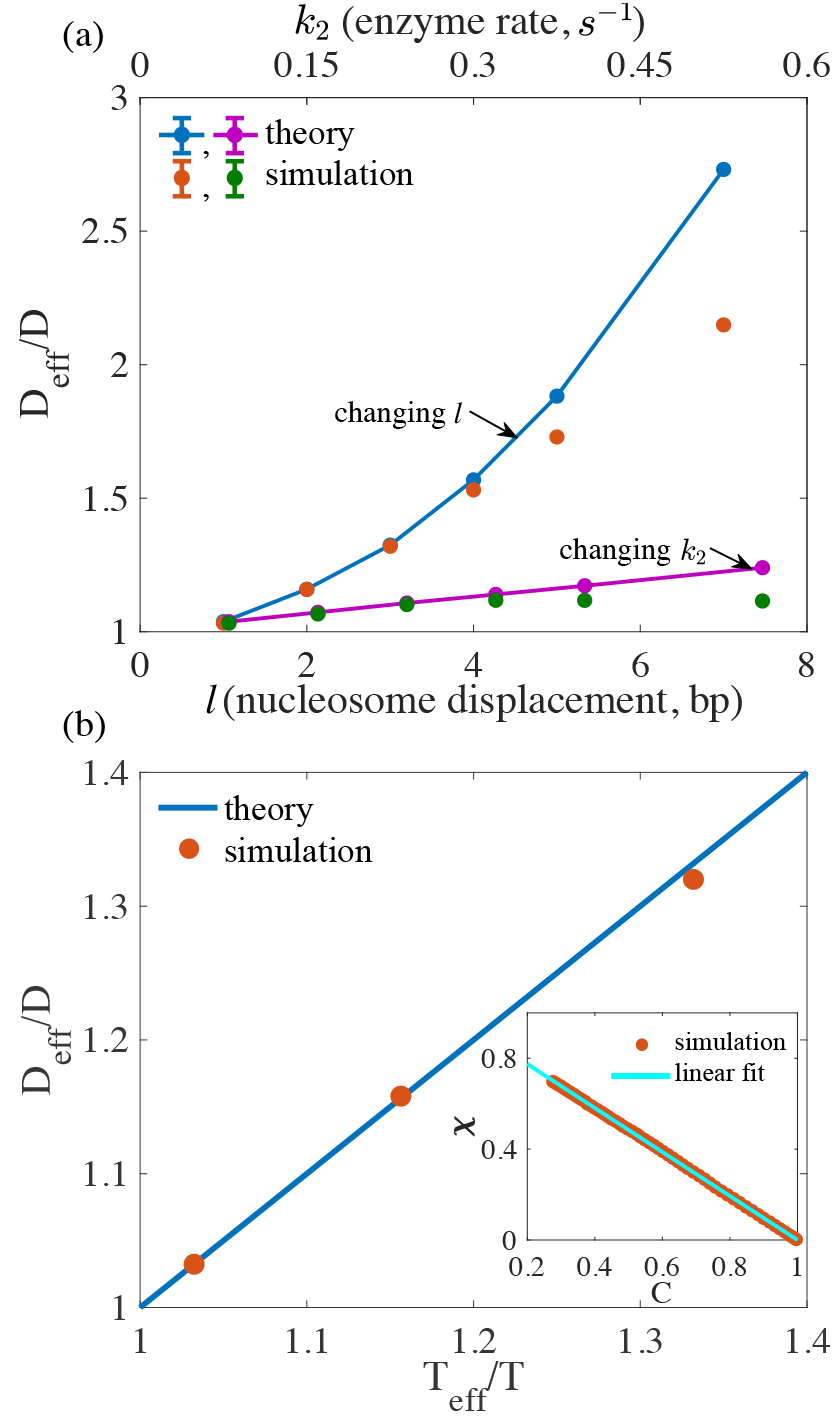
Comparison between simulated and theoretical values of *D*_eff_ and *T*_eff_ for type two enzymes. (a) Dependence of *D*_eff_ on the enzyme rate *k*_2_ and the nucleosome displacement *l* per enzymatic step. (b) *T*_eff_ determined as the fluctuation-dissipation ratio at *l* = 1, 2, 3 bp and *k*_2_ = 0.08 s^−1^. Error bars measured as standard deviation of the mean are comparable to the size of the symbols.

Type two enzymes significantly impact steady-state nucleosome positioning, and *g*(*r*) exhibits substantially more peaks even at *l* = 1 bp (empty circles in Figure 5). They lead to the formation of nucleosome clusters, which will further precipitate into a single liquid phase at large *l*. Due to the constraints in one dimension, formation of nucleosome clusters increases the timescale needed to reach steady state distribution and presents a challenge for numerical simulations. We therefore limited our comparison between theory and simulation to small *l* values. For *l* = 1 and 2 bp, equilibrium *g*(*r*) determined from the effective models (cyan lines) are again in good agreement with steady state results. For *l* = 3 bp, we carried out stochastic simulations for both the full kinetic model (empty circles) and the effective model (red line) for 1.9 × 10^8^ s to calculate the *g*(*r*). These simulations are not long enough to reach steady state or equilibrium distributions, as the *g*(*r*) differ from the true equilibrium result determined from the partition function. However, the simulation results are in excellent agreement with each other, supporting the accuracy of the effective model in predicting time-dependent quantities.

**FIG. 5.**
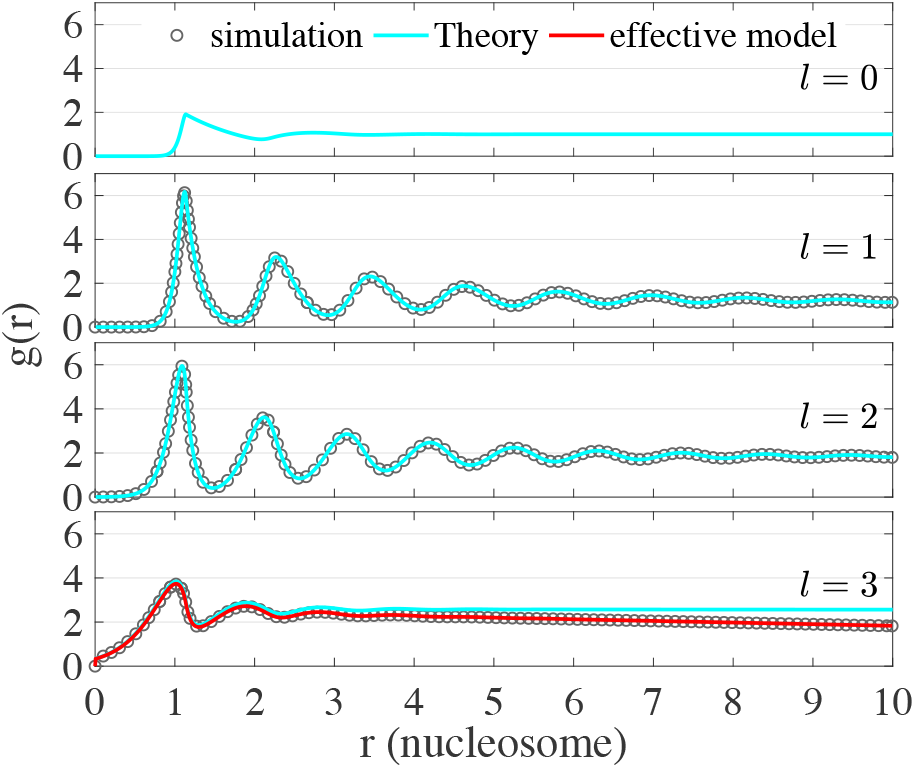
Comparison between radial distribution functions obtained from non-equilibrium simulations (empty circles) and from theoretical predictions of the effective equilibrium model (cyan lines) for type two enzymes with various step sizes. The red line for *l* = 3 bp was obtained from numerical simulations of the effective equilibrium model.

Our study provides a theoretical understanding for the interplay between remodeling enzymes and intrinsic signals in positioning nucleosomes. In particular, enzymes such as SWI/SNF that act on single nucleosome substrates do not alter the energy landscape dictated by the DNA sequence and nucleosome-nucleosome interactions. They may, in fact, reinforce the effect of these intrinsic signals by accelerating the rate for the system to approach equilibrium. On the other hand, type two enzymes can overwrite the intrinsic signals with effective interactions, leading to steady state distributions that differ from the equilibrium one [34]. The theoretical framework laid down here also provides support for using effective equilibrium models derived from the maximum entropy principle to study non-equilibrium systems [35–38].

We thank Jianshu Cao and Mehran Kardar for helpful discussions. This work was supported by startup funds from the Department of Chemistry at the Massachusetts Institute of Technology. Z.J. acknowledges the Amy Lin Shen Fellowship for financial support.

