## Supporting information for "Theory of active chromatin remodeling"

### I. DERIVATION OF EFFECTIVE EQUILIBRIUM MODELS

Here, we provide detailed derivations for the effective equilibrium models presented in the main text (Eqs. 5-7). First, we combine the non-equilibrium operators introduced in Eqs. 3 and 4 of the main text into the following compact form.

$$\hat{L}_{\rm NE}\Psi(\mathbf{x},t)) = \frac{k}{2}\sum_i [C_{i,i+1}\Psi(\mathbf{x}^i_{-l},t) + C_{i,i-1}\Psi(\mathbf{x}^i_{l},t) - (C_{i,i+1}+C_{i,i-1})\Psi(\mathbf{x},t)] \tag{S1}$$

where $\mathbf{x}^i_{\pm l} = \{\cdots, x_i \pm l, \cdots\}$. The neighboring function $C_{i,i\pm1}$ is defined as

$$C_{i,i\pm1} = \begin{cases} \text{type one enzymes :} & \begin{cases} 1 & 0 < |x_{i,i\pm1}| \\ 0 & \text{otherwise} \end{cases} \\ \text{type two enzymes :} & \begin{cases} 1 & 0 < |x_{i,i\pm1}| \leq \Delta x_{\rm max} \\ 0 & \text{otherwise} \end{cases} \end{cases} \tag{S2}$$

Here, $\Delta x_{\rm max} = 332$ bp and $x_{i,i+1} := x_{i+1} - x_i$. We note that the expression of $C_{i,i\pm1}$ for type two enzymes is identical to the one introduced in the main text. Since type one enzymes only operate on single-nucleosome substrates, their activity is not influenced by neighboring nucleosomes, and $C_{i,i\pm1}$ always equal to 1 unless the two nucleosomes are on top of each other. Replacing $C_{i,i\pm1}$ with 1, Eq. S1 reduces to Eq. 3 of the main text for type one enzymes.

Next, expanding Eq.(S1) in terms of $l$ up to the quadratic order and combining with $\hat{L}_{\rm FP}$ lead to the following equations

$$\begin{aligned}
&(\hat{L}_{\rm FP} + \hat{L}_{\rm NE})\Psi(\mathbf{x},t) \\
&= \sum_i [D\cdot\frac{\partial^2\Psi}{\partial x_i^2} + \frac{\partial}{\partial x_i}(D\beta\cdot\Psi\cdot\frac{\partial U}{\partial x_i})] + \frac{1}{2}k\sum_i\sum_{j=i\pm1} C_{ij}\cdot[\frac{1}{2}l^2\frac{\partial^2\Psi}{\partial x_i^2} - l\frac{\partial\Psi}{\partial x_i}\hat{x}_{ij}\cdot\hat{x}_i] \\
&= \sum_i [(D + \frac{1}{4}kl^2\sum_{j=i\pm1}C_{ij})\cdot\frac{\partial^2\Psi}{\partial x_i^2} + \frac{\partial}{\partial x_i}(D\beta\cdot\Psi\cdot\frac{\partial U}{\partial x_i}) - \frac{1}{2}kl\frac{\partial\Psi}{\partial x_i}\sum_{j=i\pm1}C_{ij}\cdot\hat{x}_{ij}\cdot\hat{x}_i] \\
&= \sum_i\{\frac{\partial}{\partial x_i}[(D + \frac{1}{4}kl^2\sum_{j=i\pm1}C_{ij})\cdot\frac{\partial\Psi}{\partial x_i}] + \frac{\partial}{\partial x_i}(D\beta\cdot\Psi\cdot\frac{\partial U}{\partial x_i})\} \\
&\quad - \sum_i\frac{\partial\Psi}{\partial x_i}[\frac{\partial}{\partial x_i}(D + \frac{1}{4}kl^2\sum_{j=i\pm1}C_{ij}) + \frac{1}{2}kl\sum_{j=i\pm1}C_{ij}\cdot\hat{x}_{ij}\cdot\hat{x}_i] \\
&= \sum_i[\frac{\partial}{\partial x_i}(D^i_{\rm eff}\cdot\frac{\partial\Psi}{\partial x_i}) + \frac{\partial}{\partial x_i}(D^i_{\rm eff}\beta^i_{\rm eff}\cdot\Psi\cdot\frac{\partial U}{\partial x_i}) + \frac{\partial}{\partial x_i}(D^i_{\rm eff}\beta^i_{\rm eff}\cdot\Psi)\cdot\frac{\partial\Lambda^i_{\rm mod}}{\partial x_i}] \\
&= \sum_i[\frac{\partial}{\partial x_i}(D^i_{\rm eff}\cdot\frac{\partial\Psi}{\partial x_i}) + \frac{\partial}{\partial x_i}(D^i_{\rm eff}\beta^i_{\rm eff}\cdot\Psi\cdot\frac{\partial U_{\rm eff}}{\partial x_i})],
\end{aligned} \tag{S3}$$

where the modified potential satisfies

$$\frac{\partial \Lambda^i_{\rm mod}}{\partial x_i} = -(D\beta)^{-1}[\frac{\partial}{\partial x_i}(D + \frac{1}{4}kl^2 \sum_{j=i\pm1} C_{ij}) + \frac{1}{2}kl \sum_{j=i\pm1} C_{ij} \cdot \hat{x}_{ij} \cdot \hat{x}_i] \tag{S4}$$

In deriving the expression for $\Lambda^i_{\rm mod}$, we assumed that $\frac{\partial^2 \Lambda^i_{\rm mod}}{\partial x_i^2} = 0$. As will be shown later on, this assumption is indeed self-consistent.

We note that for type one enzymes the modified potential is precisely zero. This is because that $(D + \frac{1}{4}kl^2 \sum_{j=i\pm1} C_{ij})$ is a constant, so the first term in square brackets of Eq. S4 is zero. Furthermore, since $\hat{x}_{i,i+1} = \frac{\vec{x}_{i+1} - \vec{x}_i}{|x_{i,i+1}|}$ is in the same direction with $\vec{x}_i$, $\hat{x}_{i,i+1} \cdot \hat{x}_i = 1$. Similarly, we have $\hat{x}_{i,i-1} \cdot \hat{x}_i = -1$. Therefore, the 2$^{\rm nd}$ term in the brackets is zero as well. With $\Lambda^i_{\rm mod} \equiv 0$, the expression for type one enzymes is significantly simplified, and one can readily derive the following expressions

$$\begin{aligned} D_{\rm eff} &= D + \frac{1}{2}kl^2 \\ T_{\rm eff} &= TD_{\rm eff}/D \\ U_{\rm eff} &= U(\mathbf{x}). \end{aligned} \tag{S5}$$

To obtain an expression of $\Lambda^i_{\rm mod}$ for type two enzymes, we integrate $\frac{\partial \Lambda^i_{\rm mod}}{\partial x_i}$ from $x_{i-1}$ to $x_i$

$$\begin{aligned}
\Lambda^i_{\rm mod} &= -(D\beta)^{-1}\{\int_{x_{i-1}}^{x_i} dx_i[\frac{\partial}{\partial x_i}(D + \frac{1}{4}kl^2 \sum_{j=i\pm1} C_{ij}) + \frac{1}{2}kl \sum_{j=i\pm1} C_{ij} \cdot \hat{x}_{ij} \cdot \hat{x}_i]\} \\
&= -(D\beta)^{-1}\{[\int_{x_{i-1}}^{x_i} dx_i \frac{\partial}{\partial x_i}(\frac{1}{4}kl^2 \sum_{j=i\pm1} C_{ij}) + \frac{1}{2}kl(\int_{x_{i+1}}^{x_i} C_{i,i+1}dx_i - \int_{x_{i-1}}^{x_i} C_{i,i-1}dx_i)] + f(x_{i-1}, x_{i+1})\} \\
&= -(D\beta)^{-1}\{\frac{1}{4}kl^2 \sum_{j=i\pm1} (C_{ij} + {\rm const.}^j) + \frac{1}{2}kl[(\int_{x_{i+1}}^{x_{i+1}-\Delta x_{\rm max}} C_{i,i+1}dx_i + \int_{x_{i+1}-\Delta x_{\rm max}}^{x_i} C_{i,i+1}dx_i) \\
&\qquad - (\int_{x_{i-1}}^{x_{i-1}+\Delta x_{\rm max}} C_{i,i-1}dx_i + \int_{x_{i-1}+\Delta x_{\rm max}}^{x_i} C_{i,i-1}dx_i)] + f(x_{i-1}, x_{i+1})\} \\
&= -(D\beta)^{-1}\{\frac{1}{4}kl^2 \sum_{j=i\pm1} (C_{ij} + {\rm const.}^j) + f(x_{i-1}, x_{i+1}) \\
&\qquad + \frac{1}{2}kl[-C_{i,i+1}x_{i,i+1} - (1 - C_{i,i+1})\Delta x_{\rm max} - C_{i,i-1}x_{i-1,i} - (1 - C_{i,i-1})\Delta x_{\rm max}]\} \\
&= (D\beta)^{-1}\frac{1}{2}kl \sum_{j=i\pm1} [-\frac{1}{2}C_{ij}l + C_{ij} \cdot |x_{ij}| + (1 - C_{ij}) \cdot \Delta x_{\rm max} + {\rm const.}^j].
\end{aligned} \tag{S6}$$

As will be shown later, the constant ${\rm const.}^j$ can be determined from the asymptotic condition of the pair-wise potential between neighboring nucleosomes. $f(x_{i-1}, x_{i+1})$ equals to

$\int_{x_{i-1}}^{x_{i+1}} dx_i(\frac{1}{2}kl \cdot C_{i,i+1} \cdot \hat{x}_{i,i+1} \cdot \hat{x}_i)$, and is a function only related to $x_{i\pm 1}$. Therefore, $f(x_{i-1}, x_{i+1})$ can be incorporated into const.$^j$.

As we can seen from Eq. S6, the modified potential $\Lambda^i_{\text{mod}}$ is a linear function of $x_i$. Therefore, our assumption on that the second derivative $\frac{\partial^2 \Lambda^i_{\text{mod}}}{\partial x_i^2} = 0$ is valid. Strictly speaking, however, $\frac{\partial^2 \Lambda^i_{\text{mod}}}{\partial x_i^2}$ diverges at the point $x_i = \Delta x_{\max}$ due to the discontinuity of $C_{ij}$. Practically, we find that this approximation has no impact on the accuracy of the theory.

Since each nucleosome $i$ will interact with both its left and right neighbors, a pair wise modified from potential can be derived following Eq. S6 as

$$\lambda_{\text{mod}}(x_{i,i+1}) = (D\beta)^{-1}\frac{1}{2}kl[-\frac{1}{2}C_{i,i+1}l + C_{i,i+1} \cdot x_{i,i+1} + (1 - C_{i,i+1}) \cdot \Delta x_{\max} + \text{const.}^i]. \quad \text{(S7)}$$

The constant const.$^i$ can be determined with the asymptotic condition $\lim_{\Delta x \to \infty} \lambda_{\text{mod}}(\Delta x) = 0$, resulting in const.$^i = -\Delta x_{\max}$.

Combining all the results above, we arrive at the following effective parameters for type two enzymes

$$D^i_{\text{eff}} = D + \frac{1}{4}kl^2 \sum_{j=i\pm 1} C_{ij} \quad \text{(S8)}$$
$$T^i_{\text{eff}} = TD^i_{\text{eff}}/D$$
$$U_{\text{eff}} = U + (D\beta)^{-1}\sum_i \frac{1}{2}kl[C_{i,i+1} \cdot (x_{i,i+1} - \Delta x_{\max}) - \frac{1}{2}C_{i,i+1}l]$$

A mean field approximation is further applied to replace $C_{ij}$ with $\overline{C} = \frac{1}{N}\sum_i \langle C_{i,i+1} \rangle$ to derive an approximate by constant effective diffusion coefficient and constant effective temperature shown in Eq. 7 of the main text.

### II. DETAILS OF KINETIC SIMULATIONS

To determine kinetic and thermodynamic quantities of the non-equilibrium system, we carried out stochastic simulations of the lattice model for nucleosome positioning using the Gillespie algorithm [1]. A 58800 bp long DNA with a total of 400 binding sites was used together with the periodic boundary condition. Only one type of enzymes is included in any given simulation in order to study their effects separately. We used a nucleosome density of 0.8 for simulations with type one enzymes and 0.5 for type two enzymes. These densities were chosen to highlight the effect of the enzymes and do not affect our conclusions.

All the simulations for type one enzymes were initialized with configurations in which the nucleosomes are placed next to each other. All the simulations for type two enzymes were initialized with configurations in which the nucleosomes are uniformly distributed along the lattice. As we performed long-timescale simulations to ensure their convergence, these initial configurations have no effect on the results presented in the main text. At any given time, a nucleosome can move from lattice site $m$ to $n = m \pm 1$ via diffusion with rate $d_{mn} = \frac{D}{(1\ \text{bp})^2} e^{\beta(U_m - U_n)/2}$. Nucleosomes can also be displaced with $l$ bp by remodeling enzymes at a rate of $k_{1/2}$ for type one and two enzymes respectively. We used $D = 1\ \text{bp}^2/\text{s}$, $k_1 = 1\ \text{s}^{-1}$ and $k_2 = 0.08\ \text{s}^{-1}$ unless otherwise specified.

From the simulated nucleosome configurations, we determined $D_{\text{eff}}$, $T_{\text{eff}}$, and $g(r)$ to validate the accuracy of effective equilibrium models as detailed below.

#### A. Diffusion coefficients

Diffusion coefficients were calculated from linear fits to the position mean square displacement

$$\left\langle \Delta x(\Delta t)^2 \right\rangle = \frac{1}{N} \sum_{i=1}^{N} \frac{1}{N_T} \sum_{j=1}^{N_T} \left\langle [x_i(t_j + \Delta t) - x_i(t_j)]^2 \right\rangle, \tag{S9}$$

where $N$ is the total number of nucleosomes and $N_T$ is the total number of initial configurations. To calculate the ensemble averages, for type one enzymes, we carried out ten independent $N_{\text{tot}}$-step-long simulations for each combination of $k_1$ and $l$ studied in the main text. A total of $N_{\text{T}} = 2 \times 10^4$ initial configurations were then selected uniformly along each trajectory. Only configurations from the first $\Delta t$ long trajectory of each segment was used to determine $\langle \Delta x(\Delta t)^2 \rangle$, and the rest was discarded. Exact values for $N_{\text{tot}}$, $\Delta t$, and the average time per step $\tau$ are provided in Table S1.

For type two enzymes, we followed essentially the same simulation protocol, except that the first $2 \times 10^8$-step-long segment of each trajectory was discarded as equilibration and $N_{\text{T}} = 8 \times 10^4$ initial configurations were uniformly selected. Simulation parameters are provided in Table S2. The same trajectories were used to determine $\overline{C} = \frac{1}{N} \sum_i \langle C_{i,i+1} \rangle$.

We hypothesized that the deviations between theoretical and simulated $D_{\text{eff}}$ observed in Figures 2(a) and 4(a) of the main text are caused by strong overlaps between neighboring nucleosomes. To validate this hypothesis, we carried out additional short-timescale simulations in which the system has not been "equilibrated" long enough for such configurations to appear. A total 40000 nucleosome binding sites were used in these simulations to ensure better statistics. Simulation parameters used to calculate mean-squared-displacement are provided in Tables S1 and S2 noted as $N_{\rm tot}^{\rm short}$, $\tau_{\rm tot}^{\rm short}$ and $\Delta t^{\rm short}$.

#### B. Effective temperature

As mentioned in the main text, we used the fluctuation-dissipation ratio to determine the effective temperature

$$\chi(t) = \frac{1}{T_{\rm eff}}\left[C(0) - C(t)\right]. \quad \text{(S10)}$$

We define

$$\chi(t) = \langle O(t) - O(0)\rangle / h \quad \text{(S11)}$$

$$C(t) = \langle O(t+t_o)O'(t_o)\rangle - \langle O(t_o)\rangle \langle O'(t_o)\rangle \quad \text{(S12)}$$

$$O(t) = \frac{1}{N}\sum_{j=1}^{N} \epsilon_j \exp[i\kappa \cdot x_j(t)] \quad \text{(S13)}$$

$$O'(t) = 2\sum_{j=1}^{N} \epsilon_j \cos[\kappa \cdot x_j(t)] \quad \text{(S14)}$$

$\epsilon_j = \pm 1$ follows the bimodal distribution with zero mean [2, 3]. The wave vector $\kappa$ was chosen as the value for the first peak of the structure factor defined as

$$S(\kappa) = \frac{1}{N}\sum_{j=1}^{N}\sum_{m=1}^{N} \langle \exp\{i\kappa \cdot [x_j(t_o) - x_m(t_o)]\}\rangle , \quad \text{(S15)}$$

which can be determined from the Fourier transform of $g(r)$. Numerical values for $\kappa$ are provided in Table S3.

We used the following protocol to calculate $C(t)$ and $\chi(t)$. First, 1000 initial configurations were uniformly sampled from a $10^{10}$-step-long trajectory initialized from the end state of $g(r)$ simulations for each parameter set. Starting from these configurations, we then carried out $3 \times 10^6$-step-long simulations to determine the ensemble averages for type one enzymes. Simulations of $2 \times 10^6$-step-long were performed for type two enzymes. The first $4 \times 10^5$ steps of each simulation were discarded as equilibration.

For $C(t)$, we generated 200 random realizations for $\epsilon_j$, and determined its final value using Eq. S12 by averaging over 1000 trajectories initialized from the different starting configurations mentioned above. For $\chi(t)$, we performed 200 independent simulations with random realizations of $\epsilon_j$ for each one of the 1000 initial configurations by adding a perturbation force $F_j(\kappa, t) = -\frac{\partial}{\partial x_j}[-hO'(t)]$ to each nucleosome. Specifically, each particle will experience a time-dependent perturbation field $-2h\epsilon_j \cos[\kappa \cdot x_j(t)]$. $h$ was chosen as 0.1 $k_B T$ to ensure the accuracy of the linear response theory. The final values for $\chi(t)$ were therefore determined from $1000 \times 200$ independent simulations.

After obtaining $C(t)$ and $\chi(t)$, we plot $\chi \sim C$ and the slope is the negative reciprocal of the system's effective temperature according to $\chi(t) = \frac{1}{T_{\text{eff}}}[C(0) - C(t)]$.

#### C. Radial distribution function

To determine the radial distribution function $g(r)$ for each parameter set, we performed 10 independent simulations of $N_g + N_e$ steps in length. The first $N_e$ steps of these simulations were discarded as equilibration. Nucleosome configurations were then collected at every 2000 steps for the $N_g$-step-long production segments. Numerical values for $N_g$ and $N_e$ are provided in Table S4.

For type two enzymes with step size $l = 3$ bp, we carried out stochastic simulations of both the non-equilibrium kinetic model and the effective equilibrium model defined in Eq. 7 of the main text. We note that $N_g$ for simulations of the effective equilibrium model differ from that of the non-equilibrium kinetic simulation. This is to ensure that the lengths in real time unit (second) are the same for these simulations.

### III. RADIAL DISTRIBUTION FUNCTION OF EFFECTIVE EQUILIBRIUM MODELS

Theoretical values for $g(r)$ shown in Figure 3 and 5 of the main text were calculated following Ref. [4]. In particular, for a system with $M$ binding sites, its grand canonical partition function $\Xi$ satisfies the following relationship

$$\Xi = \mathbf{e}\mathbf{W}^M\mathbf{e}^+ \tag{S16}$$

where transition probability matrix $\mathbf{W}$ is defined as

$$\begin{aligned} w_{1j} &= yq_j, \;\; j = 1 \text{ to } n_0 \\ w_{j+1,j} &= 1, \;\; j = 1 \text{ to } n_0 - 1 \\ w_{n_0,n_0} &= 1, \\ w_{i,j} &= 0, \quad \text{all other } i, j \end{aligned} \tag{S17}$$

The Boltzmann factors $q_n$ is defined as

$$q_n = \begin{cases} e^{-v(n)/k_BT} & 0 < n < n_0 \\ 1 & n \geq n_0 \end{cases} \tag{S18}$$

where $v(n)$ is the pair-wise potential between neighboring nucleosomes, and $n_0 = 294, 441$ bp for type one and two enzymes respectively. $y = \delta e^{\mu/k_BT}/A$, where $A^{-1}$ is the integral over momentum space in partition function and $\delta^{-1} = 147$ bp. $\mathbf{e}, \mathbf{e}^+$ describe the boundary conditions of lattice site occupation.

From the partition function, the system density $\rho$ and radial distribution function $g(r)$ can be determined as

$$\begin{aligned} \rho &= \mathbf{e_1} \mathbf{W}^M \mathbf{e_1^+} / (\delta \lambda_1^M) \\ g(r = n) &= \mathbf{e_1} \mathbf{W}^n \mathbf{e_1^+} / (\delta \rho \lambda_1^n) \end{aligned} \tag{S19}$$

where $\mathbf{e_1} = (1, 0, ..., 0)$, representing the occupation of the previous site ($\mathbf{e_1}^+$ for the following site) and $\lambda_1$ is the maximum eigenvalue of matrix $\mathbf{W}$.

Results shown in the main text are determined with $M = 2 \times 10^4$ and $4 \times 10^5$ for type one and type two enzymes, respectively. A larger value of $M$ for type two enzymes was used since the effective potential is more negative than the original soft core potential.

[1] D. T. Gillespie, J. Phys. Chem. **81**, 2340 (1977).

[2] L. Berthier and J. L. Barrat, J. Chem. Phys. **116**, 6228 (2002), 0111312 [cond-mat].

[3] D. Loi, S. Mossa, and L. F. Cugliandolo, Soft Matter **7**, 10193 (2011), 1105.0806.

[4] D. Poland, J. Stat. Phys. **5**, 159 (1972).

| $k_1(\mathrm{s}^{-1})$ | $l(\mathrm{bp})$ | $N_{\mathrm{tot}}$ | $\tau(\mathrm{s})$ | $\Delta t(\mathrm{s})$ | $N_{\mathrm{tot}}^{\mathrm{short}}$ | $\tau^{\mathrm{short}}(\mathrm{s})$ | $\Delta t^{\mathrm{short}}(\mathrm{s})$ |
|---|---|---|---|---|---|---|---|
| 1 | 1 | $4 \times 10^8$ | $1.04 \times 10^{-3}$ | $7.8 \times 10^{-3}$ | $6 \times 10^5$ | $1.0 \times 10^{-5}$ | $1.95 \times 10^{-4}$ |
| 1 | 2 | $4 \times 10^8$ | $1.04 \times 10^{-3}$ | $7.8 \times 10^{-3}$ | $6 \times 10^5$ | $1.0 \times 10^{-5}$ | $1.95 \times 10^{-4}$ |
| 1 | 3 | $4 \times 10^8$ | $1.04 \times 10^{-3}$ | $7.8 \times 10^{-3}$ | $6 \times 10^5$ | $1.0 \times 10^{-5}$ | $1.95 \times 10^{-4}$ |
| 1 | 5 | $4 \times 10^8$ | $1.05 \times 10^{-3}$ | $7.8 \times 10^{-3}$ | $6 \times 10^5$ | $1.0 \times 10^{-5}$ | $1.95 \times 10^{-4}$ |
| 1 | 7 | $4 \times 10^8$ | $1.05 \times 10^{-3}$ | $7.8 \times 10^{-3}$ | $6 \times 10^5$ | $1.0 \times 10^{-5}$ | $1.95 \times 10^{-4}$ |
| 1 | 10 | $4 \times 10^8$ | $1.06 \times 10^{-3}$ | $7.8 \times 10^{-3}$ | $6 \times 10^5$ | $1.0 \times 10^{-5}$ | $1.95 \times 10^{-4}$ |
| 1 | 15 | $4 \times 10^8$ | $1.07 \times 10^{-3}$ | $7.8 \times 10^{-3}$ | $6 \times 10^5$ | $1.0 \times 10^{-5}$ | $1.95 \times 10^{-4}$ |
| 1 | 20 | $4 \times 10^8$ | $1.08 \times 10^{-3}$ | $7.8 \times 10^{-3}$ | $6 \times 10^5$ | $1.0 \times 10^{-5}$ | $1.95 \times 10^{-4}$ |
| 2 | 1 | $4 \times 10^8$ | $7.81 \times 10^{-4}$ | $7.8 \times 10^{-3}$ | N/A | N/A | N/A |
| 3 | 1 | $4 \times 10^8$ | $6.25 \times 10^{-4}$ | $7.8 \times 10^{-3}$ | N/A | N/A | N/A |
| 5 | 1 | $4 \times 10^8$ | $4.47 \times 10^{-4}$ | $3.9 \times 10^{-3}$ | N/A | N/A | N/A |
| 7 | 1 | $4 \times 10^8$ | $3.48 \times 10^{-4}$ | $1.95 \times 10^{-3}$ | N/A | N/A | N/A |
| 10 | 1 | $4 \times 10^8$ | $2.61 \times 10^{-4}$ | $1.95 \times 10^{-3}$ | N/A | N/A | N/A |
| 15 | 1 | $4 \times 10^8$ | $1.84 \times 10^{-4}$ | $1.95 \times 10^{-3}$ | N/A | N/A | N/A |
| 20 | 1 | $4 \times 10^8$ | $1.42 \times 10^{-4}$ | $1.95 \times 10^{-4}$ | N/A | N/A | N/A |
| 30 | 1 | $4 \times 10^8$ | $9.80 \times 10^{-5}$ | $1.95 \times 10^{-4}$ | N/A | N/A | N/A |
| 100 | 1 | $4 \times 10^8$ | $3.08 \times 10^{-5}$ | $7.8 \times 10^{-5}$ | N/A | N/A | N/A |
| 400 | 1 | $4 \times 10^8$ | $7.82 \times 10^{-6}$ | $1.17 \times 10^{-5}$ | N/A | N/A | N/A |

TABLE S1. Simulation parameters used to calculate the effective diffusion coefficient $D_{\mathrm{eff}}$ for type one enzymes.

| $k_2(\mathrm{s}^{-1})$ | $l(\mathrm{bp})$ | $N_{\mathrm{tot}}$ | $\tau(\mathrm{s})$ | $\Delta t(\mathrm{s})$ | $N_{\mathrm{tot}}^{\mathrm{short}}$ | $\tau^{\mathrm{short}}(\mathrm{s})$ | $\Delta t^{\mathrm{short}}(\mathrm{s})$ |
|---|---|---|---|---|---|---|---|
| 0.08 | 1 | $10^9$ | $2.41 \times 10^{-3}$ | $1.95 \times 10^{-2}$ | $6 \times 10^6$ | $2.4 \times 10^{-5}$ | $3.9 \times 10^{-4}$ |
| 0.08 | 2 | $10^9$ | $2.41 \times 10^{-3}$ | $1.95 \times 10^{-2}$ | $6 \times 10^6$ | $2.4 \times 10^{-5}$ | $3.9 \times 10^{-4}$ |
| 0.08 | 3 | $10^9$ | $2.42 \times 10^{-3}$ | $1.95 \times 10^{-2}$ | $6 \times 10^6$ | $2.4 \times 10^{-5}$ | $3.9 \times 10^{-4}$ |
| 0.08 | 4 | $10^9$ | $2.44 \times 10^{-3}$ | $1.95 \times 10^{-2}$ | $6 \times 10^6$ | $2.4 \times 10^{-5}$ | $3.9 \times 10^{-4}$ |
| 0.08 | 5 | $10^9$ | $2.49 \times 10^{-3}$ | $1.95 \times 10^{-2}$ | $6 \times 10^6$ | $2.4 \times 10^{-5}$ | $3.9 \times 10^{-4}$ |
| 0.08 | 7 | $10^9$ | $2.55 \times 10^{-3}$ | $1.95 \times 10^{-2}$ | $6 \times 10^6$ | $2.4 \times 10^{-5}$ | $3.9 \times 10^{-4}$ |
| 0.08 | 10 | N/A | N/A | N/A | $6 \times 10^6$ | $2.4 \times 10^{-5}$ | $3.9 \times 10^{-4}$ |
| 0.08 | 15 | N/A | N/A | N/A | $6 \times 10^6$ | $2.4 \times 10^{-5}$ | $3.9 \times 10^{-4}$ |
| 0.08 | 20 | N/A | N/A | N/A | $6 \times 10^6$ | $2.4 \times 10^{-5}$ | $3.9 \times 10^{-4}$ |
| 0.16 | 1 | $10^9$ | $2.33 \times 10^{-3}$ | $1.95 \times 10^{-2}$ | $6 \times 10^6$ | $2.3 \times 10^{-5}$ | $3.9 \times 10^{-4}$ |
| 0.24 | 1 | $10^9$ | $2.26 \times 10^{-3}$ | $1.95 \times 10^{-2}$ | $6 \times 10^6$ | $2.2 \times 10^{-5}$ | $3.9 \times 10^{-4}$ |
| 0.32 | 1 | $10^9$ | $2.23 \times 10^{-3}$ | $1.95 \times 10^{-2}$ | N/A | N/A | N/A |
| 0.40 | 1 | $10^9$ | $2.23 \times 10^{-3}$ | $1.95 \times 10^{-2}$ | $6 \times 10^6$ | $2.2 \times 10^{-5}$ | $3.9 \times 10^{-4}$ |
| 0.56 | 1 | $10^9$ | $2.23 \times 10^{-3}$ | $1.95 \times 10^{-2}$ | $6 \times 10^6$ | $2.0 \times 10^{-5}$ | $3.9 \times 10^{-4}$ |
| 0.8 | 1 | N/A | N/A | N/A | $6 \times 10^6$ | $1.8 \times 10^{-5}$ | $3.9 \times 10^{-4}$ |
| 1.2 | 1 | N/A | N/A | N/A | $6 \times 10^6$ | $1.6 \times 10^{-5}$ | $3.9 \times 10^{-4}$ |
| 1.6 | 1 | N/A | N/A | N/A | $6 \times 10^6$ | $1.6 \times 10^{-5}$ | $3.9 \times 10^{-4}$ |
| 8 | 1 | N/A | N/A | N/A | $6 \times 10^6$ | $5.2 \times 10^{-6}$ | $7.8 \times 10^{-5}$ |
| 32 | 1 | N/A | N/A | N/A | $6 \times 10^6$ | $1.5 \times 10^{-6}$ | $3.9 \times 10^{-5}$ |

TABLE S2. Simulation parameters used to calculate the effective diffusion coefficient $D_{\mathrm{eff}}$ for type two enzymes.

| | $\kappa$ |
|---|---|
| Type one enzymes | 0.035904 |
| Type two enzymes, $l = 1$ | 0.035904 |
| Type two enzymes, $l = 2$ | 0.041033 |
| Type two enzymes, $l = 3$ | 0.046162 |

TABLE S3. Numerical values for the wave vector $\kappa$ used for the calculations of $T_{\text{eff}}$.

| | Enzyme step length ($l$) | $N_g$ | $N_e$ |
|---|---|---|---|
| Type one enzymes | $l = 1, 2, 5$ | $4e^{10}$ | $4e^{10}$ |
| Type two enzymes | 1 | $4e^{10}$ | $4.6e^{11}$ |
| | 2 | $4e^{10}$ | $2.4e^{12}$ |
| | 3 | $4e^{10}$ | $4e^{8}$ |
| | 3, effective model | $5.08e^{10}$ | $5.08e^{10}$ |

TABLE S4. Simulation parameters used to calculate the radial distribution function $g(r)$.

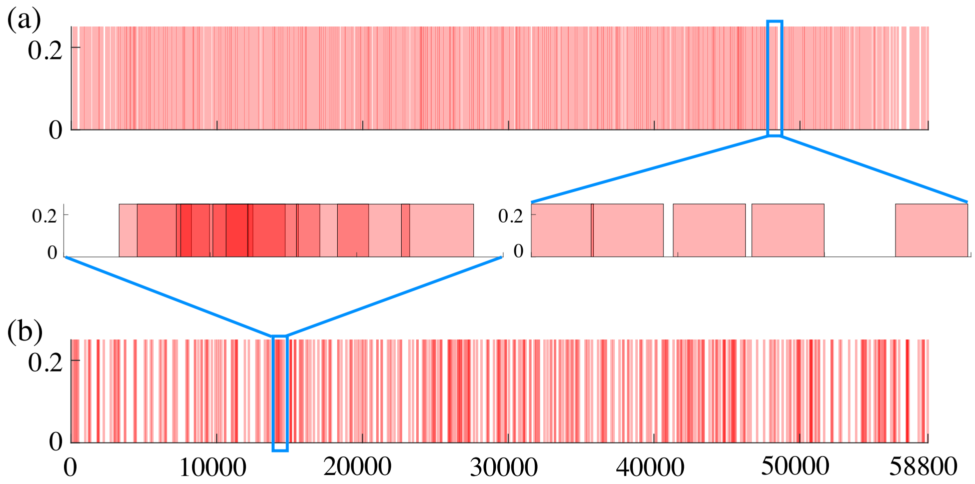


FIG. S1. Example nucleosome configurations obtained from kinetic simulations with type one enzymes of step size $l = 1$ bp (a) and 20 bp (b).

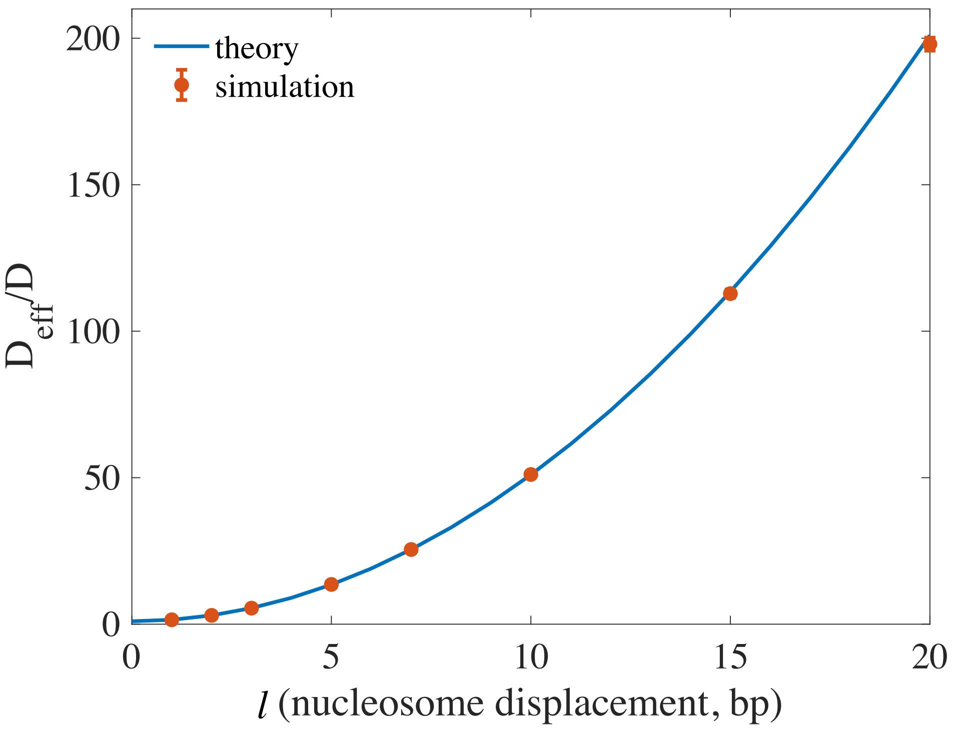


FIG. S2. Effective diffusion coefficients for type one enzymes with nucleosome displacement size $l$ obtained from short simulations, in which the system has not been evolved long enough to explore configurations with strong nucleosome overlaps.

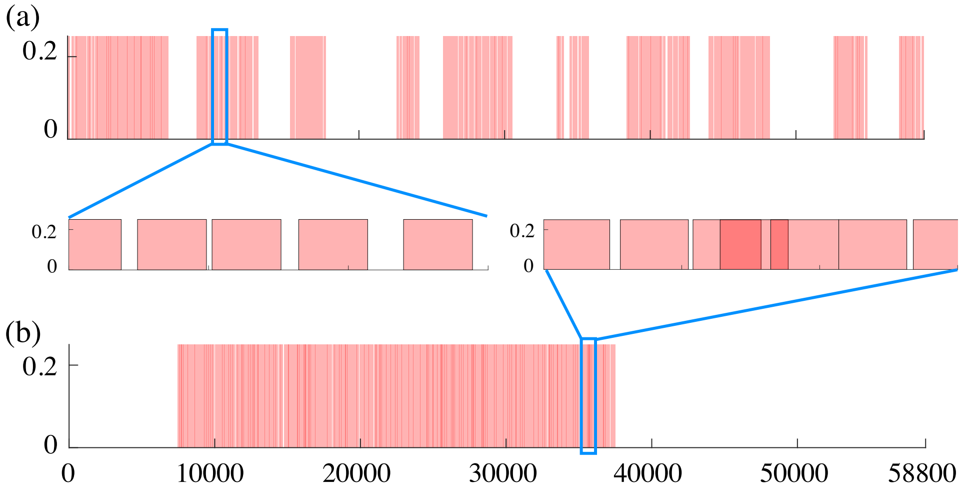


FIG. S3. Example nucleosome configurations obtained from kinetic simulations with type two enzymes of step size $l = 1$ (a) and $l = 2$ bp (b).

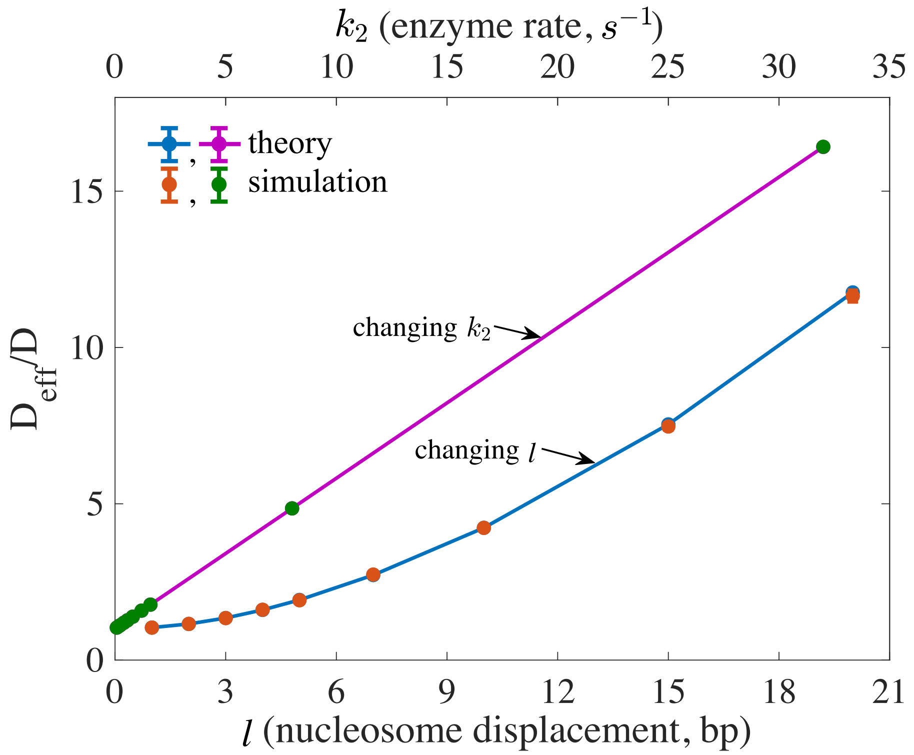


FIG. S4. Effective diffusion coefficients for type two enzymes with different nucleosome displacement size $l$ and remodeling rates $k_2$ obtained from short simulations, in which the system has not been evolved long enough to explore configurations with strong nucleosome overlaps.
